# Gene traffic mediated by transposable elements shaped the dynamic evolution of ancient sex chromosomes of varanid lizard

**DOI:** 10.1101/2025.04.01.646716

**Authors:** Zexian Zhu, Jason Dobry, Erik Wapstra, Qi Zhou, Tariq Ezaz

## Abstract

Lizards exhibit rapid turnovers and a much greater diversity of sex determination mechanisms compared to birds and eutherians. This makes the conserved ZW sex chromosomes of anguimorph lizards originated over 115 million years ago a seeming exception. We recently discovered in an anguimorph lizard *Varanus acanthurus* (*Vac*) that its entire W chromosome (chrW), but not chrZ is homologous to part of the chr2 by cytogenetic mapping, suggesting a complex history of its sex chromosome evolution yet to be elucidated. To address this, we assembled a chromosome-level genome, and provided evidence that the *Vac* sex chromosome pair had undergone at least three times of recombination suppression, forming a similar pattern of ‘evolutionary strata’ to that of birds or mammals. We identified the putative sex-determining genes in the oldest evolutionary stratum that had first lost recombination. Comparison to other lizard genomes dated the stepwise propagation of specific retrotransposon subfamilies shared by chrW and chr2 to the varanid ancestor. These retrotransposons are also enriched near the duplicated genes shared by the two chromosomes and probably mediated the recruitment of many autosomal genes that rejuvenated the degenerating chrW, including members of a large vomeronasal chemosensory receptor gene family *V2R*. Our results challenge the canonical model of sex chromosome evolution, and suggest that the W or Y chromosome as a refugium of repetitive elements, may recurrently recruit short-lived functional genes responsible for sexual dimorphisms during its long-term course of degeneration.

## Introduction

Lizards harbor a rich diversity of sex determination (SD) mechanisms that is unparalleled by other land vertebrates (Ezaz et al. 2009; Mezzasalma et al. 2021), providing great opportunities for answering fundamental questions like why and how these different mechanisms emerge and transit to each other, and how do sex chromosomes subsequently evolve. Such a diversity has long been recognized by early systematic cytogenetic reports of male heterogametic (male XY, female XX, like mammals), female heterogametic (male ZZ, female ZW, like birds), multiple (e.g., X_1_X_2_Y in *Calyptommatus* lizard due to autosome fusion with the Y chromosome (Yonenaga-Yassuda et al. 2005)) sex chromosomes in different species (Bull 1980), and temperature sex determination mechanism (TSD) that co-exists with one of the genetic sex determination (GSD) mechanisms in some species (e.g., spotted snow skink (Hill et al. 2018)). Another aspect of the diversity is the diversified rate of SD turnovers between different lizard lineages. A classic RAD-seq study identified between 17 to 25 SD transitions from 12 representative gecko species (Gamble et al. 2015). By contrast, a broad qPCR study characterized an extraordinarily conserved ZW sex chromosome pair shared by almost all studied Anguimorpha (Gila monster, beaded lizards, alligator lizards and varanids) species except for slowworm (Rovatsos et al. 2019), and the very recently reported crocodile lizards (Pinto et al. 2024). The latter strong evolutionary stasis seems to support the ‘evolutionary trap’ hypothesis (Bull and Charnov 1985; Pokorná and Kratochvíl 2009) that heteromorphic sex chromosomes resulting from recombination suppression precludes transitions into other GSD systems, as suggested by sex chromosomes of birds and mammals.

However, the fine and complex course of lizard sex chromosome evolution can be concealed due to the limited coverage and resolution of cytogenetic/qPCR/Illumina based methods, particularly the absence of abundant sex-specific chrY or chrW sequences. Although a recent burst of chromosome level lizard genomes produced by long-reads has characterized the euchromatic chrX or chrZ (Westfall et al. 2021; Geneva et al. 2022; Koochekian et al. 2022; Davalos-Dehullu et al. 2023; Leitão et al. 2023; Webster et al. 2024), little is known about their often-heterochromatic homologs chrY or chrW that likely encompass the upstream SD genes. We previously identified the candidate SD genes of an Anguimorpha species *Varanus acanthurus* (*Vac)* using an Illumina-based scaffold-level genome (Zhu et al. 2022). Based on this draft genome, we discovered a curious homology using fluorescence *in situ* hybridization between the entire chrW, but not chrZ, and the tip of autosome chr2 (**Figure 1a**, termed chr1 in the previous studies (Kinga and King 1975; Johnson et al. 2016; Iannucci et al. 2019), **Supplementary Note**) (Dobry et al. 2025), using random probes specifically targeting the chrW. This new finding questioned the reported simple history of deep evolutionarily static varanid sex chromosomes that originated over 115 million years ago (Rovatsos et al. 2019).

**Figure 1.**
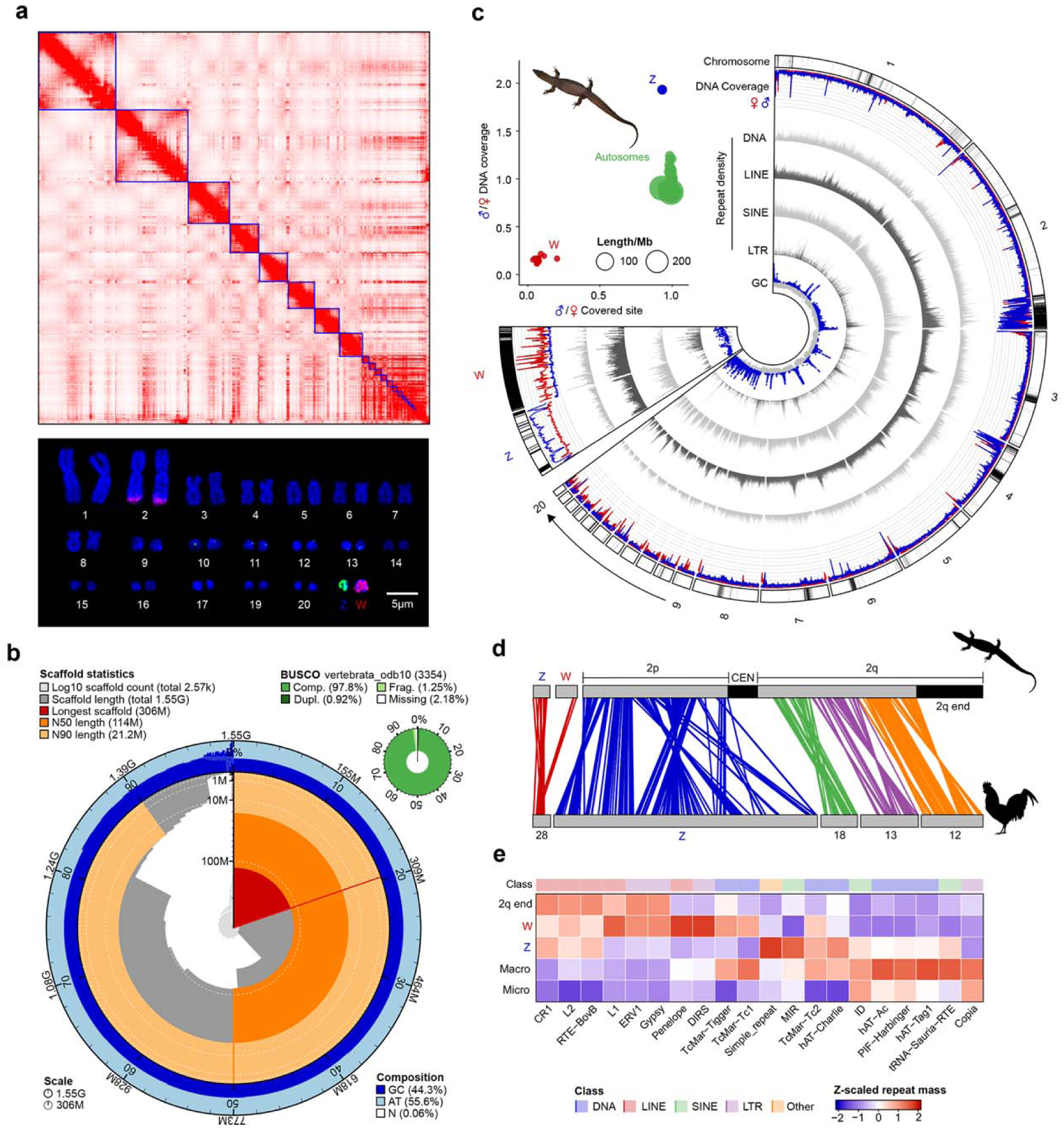
A new chromosome level genome and sex chromosomes of *Varanus acanthurus* **a)** Hi-C contact map of the *Varanus acanthurus* (*Vac*) genome. The blue square indicates an intact chromosome with preferential interactions (red dots) within rather than between each chromosome. We show the karyotype of *Vac* below, and the DNA-FISH results using the probes designed against the sequences of chrW (pink) and chrZ (green) (Dobry et al. 2025). **b)** The snail plot shows the major statistics of genome assembly including the number and length of the scaffolds, the length of the longest scaffold, the scaffold N50 and N90 values and the BUSCO score based on the vertebrate protein (vertebrate_odb10) dataset. **c)** The bubble plot shows the identification of sex chromosomes, with the bubble size scaled to the scaffold size. The chrW-linked scaffolds show a male-to-female ratio of mappable sites close to zero, whereas this ratio is around 1 for chrZ and autosomes. In addition, chrZ shows nearly a two-fold male-to-female read coverage ratio compared to the autosomes, while the ratio for the chrW remains close to zero. The circos plot shows from the outer track to inner ones: the density of overall annotated repetitive elements; DNA read coverage of male (blue) and female (red) individuals; density of four classes of repetitive elements; and the GC content. All values are calculated in 100 kb windows. **d)** Homology between the sex chromosomes of *Vac* and autosomes of chicken, with each line representing one orthologous gene pair between the two species. **e)** The heatmap shows the row-scaled TE content divided by class and family across chromosomes (Macro/micro refers to mean values calculated from macro/micro chromosomes).

Three hypothetical scenarios that are not mutually exclusive may explain the cytogenetic homology: first, the chrW has acquired duplicated sequence segments from chr2 after it stopped recombination with the chrZ. Second, the chr2 may have acquired duplicated sequences specifically from the chrW. And the last, certain repetitive sequences have accumulated at both the chr2 tip and the chrW but nowhere else in the genome, possibly because both regions have low or no recombination. Any of these scenarios may result in gene movements between sex chromosomes and autosomes that undergo very different evolutionary and molecular processes. Previous studies reported gene movements between the chrX and autosomes (termed ‘gene traffic’) in mammals (Emerson et al. 2004) and *Drosophila* (Betrán et al. 2002; Vibranovski et al. 2009). An excess of genes that move out of the chrX, producing a paucity of male-biased X-linked genes relative to autosomes can be the result of sexual antagonistic selection (Wu and Xu 2003) that predicts the chrX will undergo demasculinization of gene expression (Sturgill et al. 2007) because it is more often inherited in the female than in male. It can be also explained by other molecular processes including the meiotic X-inactivation (Turner 2007), and dosage compensation (Bachtrog et al. 2010) that specifically affect the sex chromosomes (Vicoso and Charlesworth 2006). Individual cases of either direction of gene movement, but not a scale as large as what we observed in *Vac* (**Figure 1a**), have also been reported on the human chrY (Hughes et al. 2015; Xu et al. 2020), the avian chrW (Xu et al. 2020), and the UV sex determination system of kelp (Lipinska et al. 2017). Therefore, the *Vac* sex chromosomes provide a model for elucidating the molecular and evolutionary mechanisms of large sequence exchanges between sex chromosomes and autosomes. To test the three mentioned hypotheses, in this work we produced a high-quality chromosome-level genome of a female *Vac* individual, and inferred the evolutionary history of its sex chromosomes, particularly the origin of the large sequence homology between the chrW and chr2.

## Results

### A high-quality genome of *Varanus acanthurus*

About 100-fold PacBio long read sequences were produced in order to acquire nearly complete genome and sex chromosome sequences. We anchored 97.3% of the genome (with an assembled size of 1.55 Gb vs. the estimated size of 1.5 Gb (Zhu et al. 2022)) into 19 autosomes and a chrZ by Hi-C chromatin contact data (**Figure 1a, Supplementary Table S1**). The high quality of the genome can be evidenced by the large contig N50 length (114 Mb), high BUSCO value (**Figure 1b**, 97.8% complete), and the annotated centromeric regions that are specifically enriched for L1 elements relative to the rest of the chromosome region (**Figure 1c, Supplementary Fig. 1**). Similar to other reptiles including birds, the 19 autosomes include 8 macrochromosomes, and 11 microchromosomes of smaller sizes (< 30Mb) but much higher gene density (42 genes per Mb vs. 21 of macrochromosomes).

We identified the chrZ of 11.8 Mb long, with an expected 2-fold higher genomic read depth in males than in females across most of the region (the sexually differentiated region, SDR) lacking homologous recombination in females. The chrZ is homologous to the microchromosome chr28 of chicken (**Figure 1d**), consistent with previous reports (Rovatsos et al. 2019; Zhu et al. 2022). One chromosome end of pseudoautosomal region (PAR) of only 0.57 Mb long shows an equal read depth between sexes and likely retains the recombination. We also identified a total of 418 scaffolds totaling 15 Mb, which can be mapped by female but not male reads, as expected for chrW sequences (See **Methods**). Besides the annotated centromeric regions, one chromosome end of chr2q and chr4, and the W-linked sequences have a significantly (*P* < 0.05, Wilcoxon test) higher overall repeat content, particularly in certain subfamilies of transposable elements (TEs, e.g., L1, Gypsy, **Figure 1e, Supplementary Fig. 2**), than the rest genome (**Figure 1c**). Although the chrW is depleted (*P* < 0.05, Wilcoxon test) for some families of DNA transposons and short interspersed nuclear elements (SINEs). This supports the idea that sex-specific chromosomes act as a refugium for recently active TEs (Peona et al. 2021) because they cannot be effectively purged by natural selection under a non-recombining environment (Charlesworth and Charlesworth 2000).

### Varanid sex chromosomes have undergone three recombination suppression events

A shared feature of amniote sex chromosomes, regardless XY or ZW systems, is that they usually underwent stepwise complete suppression of recombination, producing a punctuated pattern of pairwise sequence divergence levels between adjacent sex-linked regions termed ‘evolutionary strata’ (Vicoso et al. 2013; Cortez et al. 2014; Zhou et al. 2014). By change-point analyses (see **Methods**), we demarcated the Z-linked SDR into three evolutionary strata (from the old to young stratum termed as S0,1,2, **Figure 2a**), by their significantly (*P* < 0.05, Wilcoxon test) and consistently sharp differences of ZW pairwise sequence identity (% of identical bases per 100kb Z-linked sequences) and Z-linked male heterozygosity (SNP density calculated by male reads aligning to the chrZ) levels compared to the adjacent strata. A significantly different level of Z-linked heterozygosity between strata can be explained by the different time span that each stratum region has experienced with reduction of effective population size due to that of recombination (Charlesworth et al. 1987). This, to our knowledge, provides the first demonstrated case of evolutionary strata in lizards.

**Figure 2.**
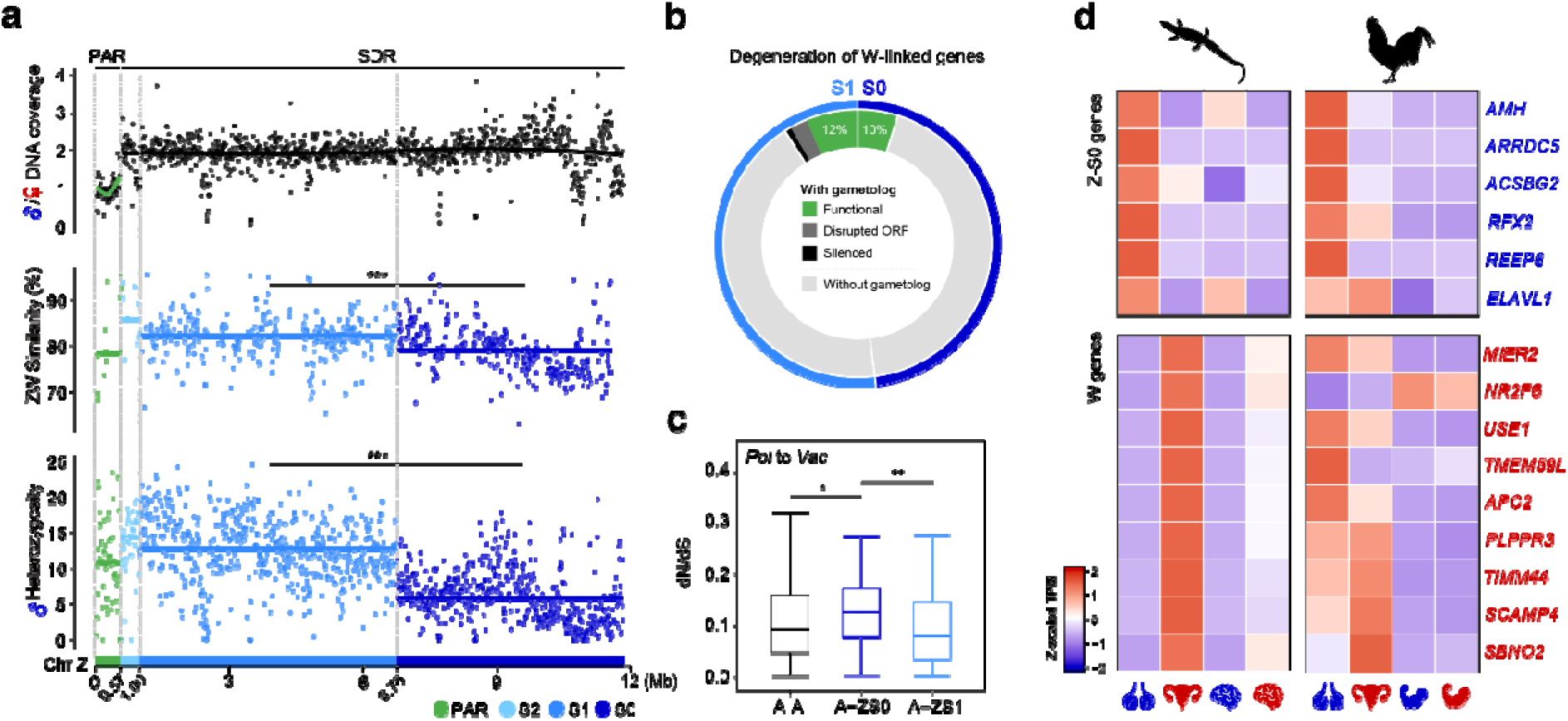
Evolutionary strata and candidate sex determining genes of *Varanus acanthurus.* **a)** *Vac* sex chromosomes consist of at least three evolutionary strata. From top to bottom, we show the male-to-female ratio of DNA read coverage; pairwise sequence identity between the chrZ and chrW; and the heterozygosity level (SNP density) calculated by male reads along the chrZ. **b)** Donut plot illustrates the degrees of degeneration of W-linked genes, divided by their residing evolutionary strata (S1 and S0). The inner ring shows the genes with gametologs present on both chrZ and chrW (‘with gametologs’), among which chrW gametologs are divided into those with intact ORFs and robust gene expression (‘functional’, green), with disrupted ORFs (grey), and with no expression (black). ‘without gametolog’ refers to genes whose chrW gametologs have been deleted. **c)** Boxplots show the pairwise dN/dS values of 1:1 orthologs between *Phrynosoma platyrhinos* (*Ppl*) and *Varanus acanthurus* (*Vac*), divided into those located on the autosomes of both species (A-A), those on the autosome in *Ppl*, but in S0 stratum of chrZ in *Vac* (A-ZS0), and those on autosome in *Ppl*, but S1 of chrZ in *Vac* (A-ZS1). **d)** The heatmap displays the z-scaled normalized expression levels (TPM) of S0 genes on the chrZ, which were reported to be involved in sex-determination pathways in other vertebrates (see **Supplementary Table S3**), along with all expressed chrW genes and their chicken orthologs across different tissues.

Due to complete lack of recombination, only 10% and 12% of the S0 and S1 W-linked genes (termed ‘gametologs’) are inferred to retain functions under our strict criterion (**Materials and Methods**). The rest W-linked gametologs have either become completely deleted compared to their Z-linked counterparts, or disrupted in the open reading frames (ORFs), or silenced in expression in all examined female tissues (**Figure 2b**). This is a conserved estimate of putatively functional genes on the chrW, as some ‘silenced’ genes are likely to be expressed in other female tissues that are not included in this study. If we count those ‘silenced’ intact ORFs, the putative functional W gametologs of *Vac* range between 21 to 30 genes, a comparable number to that of human chrY. This translates to an average gene loss rate of between 2.9 to 4.54 (0.53% to 0.87% of the) genes per million years on the chrW, a comparable rate to that of human chrY (0.5% per million years (Krasovec et al. 2018)). Interestingly, the resultant hemizygous Z-linked gametologs within the older S0, but not S1 and S2 show a significantly faster evolution rate (Wilcoxon test, *P* < 0.05), measured by pairwise nonsynonymous vs. synonymous substitution rate (dN/dS ratio) to the autosomal orthologs in the desert horned lizard (*Phrynosoma platyrhinos*, *Ppl*) (Koochekian et al. 2022), than autosomal genes, indicating a similar faster-Z effect (Charlesworth et al. 1987; Mank et al. 2010) reported in birds (Wang et al. 2014) and some other female heterogametic sex systems (Vicoso et al. 2013; Sackton et al. 2014) (**Figure 2c**).

We previously inferred the candidate sex determining gene of *Vac* to be the Z-linked *Amh*, which has a high expression in the testis and male brain, but almost no expression in the ovary and female brain (Zhu et al. 2022). Here we confirmed that *Amh* is located in the S0 region of chrZ, and we have not found its homolog on the chrW (**Figure 2d**). This is consistent with the evolutionary hypothesis that recombination was first lost at the region encompassing the sex determining gene, as demonstrated in birds and mammals (Lahn and Page 1999; Zhou et al. 2014). While *Amh* is probably the candidate Z-linked male-determining gene given its highly conserved function across vertebrates in testis development (Adolfi et al. 2019), we found few expressed genes located within the S0 region of chrW except for one gene *TIMM44*. This gene is essential for mitochondrial functions (Bonora et al. 2006), but has not been reported to be participating in any sex determination process in other vertebrate species. Its chicken ortholog has a gonad biased expression, and its *Vac* Z-linked copy has a biased expression in the male brain (**Supplementary Fig. 3**), and the W-linked copy has a biased expression in the ovary (**Figure 2d**). Whether this gene is indeed a female determining gene of *Vac* requires future studies, particularly using the embryonic gonad samples when sex is determined.

### Stepwise propagation of transposable elements in varanids

An enrichment of TEs at both chrW and the end of chr2 (termed ‘2q-end’ hereafter, region defined by the sharp change of overall repeat content along chr2, **Methods**) (**Figure 1c, Supplementary Fig. 1**) supports one of our hypotheses that the reported cytogenetic homology is contributed by shared accumulation of TEs in both regions. We next ask when did such a TE burst occur in both regions and what are the specific TEs that accumulated. To address this, we first compared the overall TE content in the homologous regions of 2q-end of *Vac*, another anguimorph lizard *Elgaria multicarinata* (*Emu*), and the Anguimorpha outgroup *Ppl*. An enrichment of TEs, but to a significantly lower degree (*P* < 0.05, Wilcoxon test), with a much shorter spanned region (23 Mb vs. 47 Mb of *Vac*), was found at the 2q-end of *Ppl* compared to the other two species. This suggests that the 2q-end was already heterochromatic, and probably has a low recombination rate in the ancestor of Anguimorpha dated 157 million years ago (**Figure 3a, Supplementary Fig. 4**). *Vac*, *Emu* and *Ppl* together exhibit a gradient of TE enrichment level and influenced region length at the 2q-end, supporting that this region experienced stepwise propagation of TEs from the ancestor of Anguimorpha to that of varanids.

**Figure 3.**
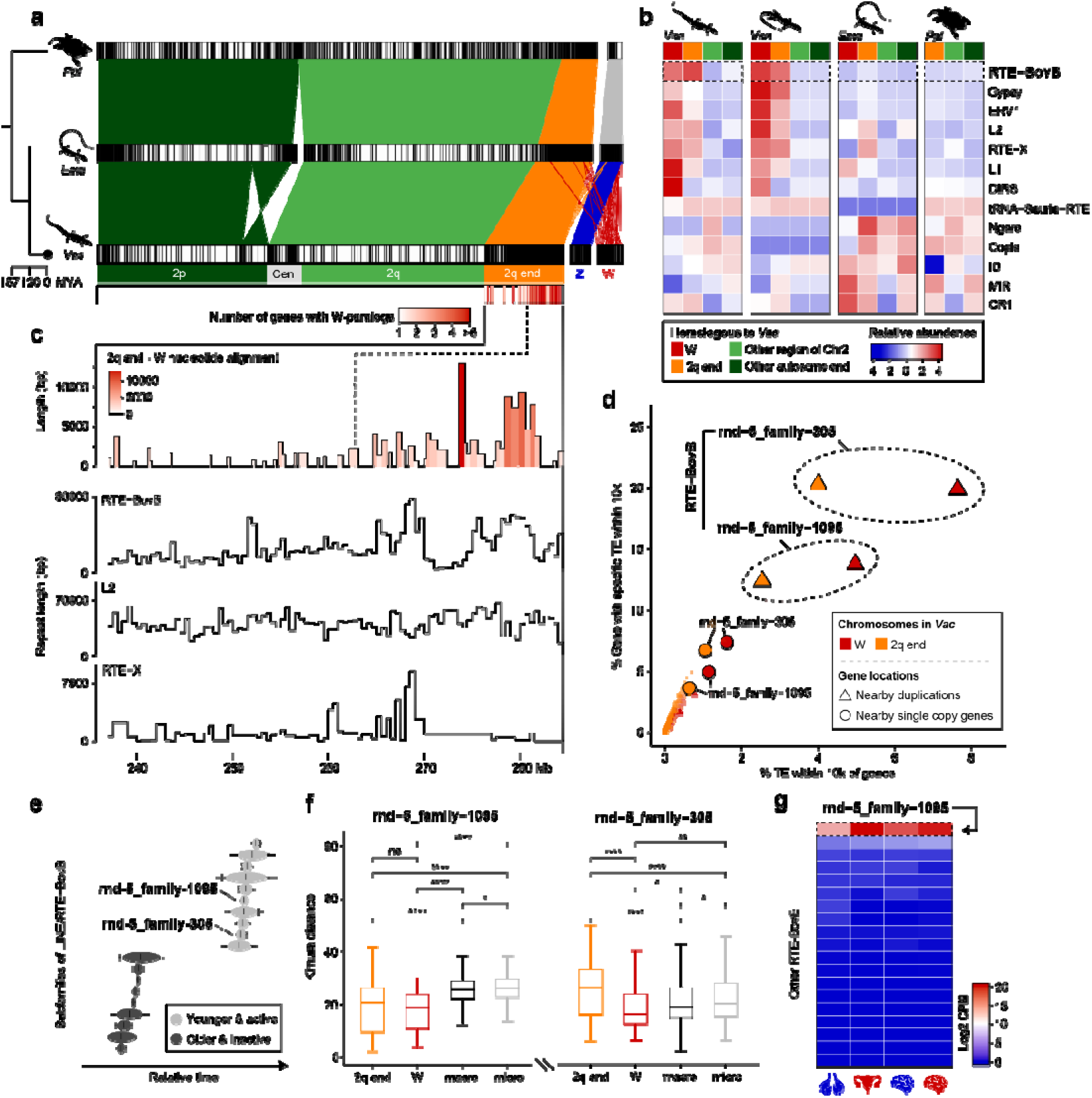
Gene duplications mediated by transposable elements between chrW and chr2q. **a)** Chromosome alignments of the sex chromosomes and chr2 in *Varanus acanthurus* (*Vac*) compared to another anguimorph lizard, *Elgaria multicarinata* (*Emu*), and the outgroup species *Phrynosoma platyrhinos* (*Ppl*). The black lines within each homologous chromosome of different species represent the density of overall TEs that shows a propagation at the 2q-end (Figure 1c). Red barcodes below indicate the number (scaled to the red color) of duplicated genes per 100 kb genomic window between chr2 and chrW. **b)** Heatmap compares the normalized repeat abundance between chrW, 2q-end, other chr2 regions, and other autosomes across representative anguimorph lizards and one outgroup lizard *Ppl*. Only LINE/RTE-BovB is enriched at the distal end of varanus lizards (*Vac* and *Vsa*) 2q-end and the chrW, compared to other genomic regions. **c)** Diagrams of the zoomed-in 2q-end region, with the first bar plot showing aligned unique region lengths with chrW. Below are line plots showing the lengths of TEs in 100kb sliding windows. The region with grey background in the first bar plot corresponds to areas with higher gene density in the barcode. **d)** Bubble plots show the enriched levels of each TE subfamily nearby single-copy genes (dots) or duplicated genes (triangles), and those of genes near each TE subfamily (**Methods**). **e)** Inferred relative ages of the subfamilies of RTE-BovB by TinT analysis, indicated by ovals and lines representing relative time points. **f)** Boxplots show the distribution of Kimura distance, another measurement of TE relative age, of all copies of rnd-5_family-1095 and rnd-5_family-305 subfamilies across different genomic regions (Macro refers to all macro chromosomes except 2q-end region and micro refers to all micro chromosomes except chrW). **g)** Heatmap shows the normalized expression levels of all RTE-BovB subfamilies across all tissues.

Particularly, we identified a long interspersed nuclear element (LINE) retrotransposon family RTE-RovB that is enriched in both chrW and 2q-end compared to the rest of genome (including chr4, whose end is also repetitive, **Figure 1c, Supplementary Fig. 5**) in both *Vac* and another varanid *V. salvator* (*Vsa*), but not in the outgroup lizard species (**Figure 3b**). While other retrotransposons, such as the long terminal repeat (LTR) families Gypsy and ERV1, show different enrichment levels between 2q-end and chrW in the two varanids. An overall 14 fold higher genome-wide percentage of RTE-RovB in varanids than their outgroup Anguimorpha species suggests an ancestral burst of this TE family in the varanid ancestor, which then preferentially inserted into the chrW and 2q-end with no or low recombination, consistent with the "TE refugium hypothesis" (Peona et al. 2021). It is also likely that the RTE-RovB elements inserted into either chromosomal region followed by transpositions onto another.

The TE burst may not be the only cause of the cytogenetic homology. And a more functionally relevant consequence is that it can mediate interchromosomal segmental duplications between the chrW and 2q-end. To investigate this, we aligned the genome sequences of both chrZ and chrW against the chr2, after masking the repeat content. The alignment of chrZ is supposed to inform us of duplications between chrW and 2q-end, followed by loss of genes on the chrW. A total of 45 paralogs of Z-linked but not W-linked genes was found on the short arm of chr2 (2p) but not in the 2q-end, 73% of which are shared with chickens (**Supplementary Fig. 6**), suggesting ancient gene duplications predating the divergence between birds and reptiles. We showed previously that the chicken homologous chromosomes chrZ and chr28 of the *Vac* chr2p and chrZ (**Figure 1d**) are derived from the same ancestral vertebrate chromosome approximated by that of amphioxus (Huang et al. 2023). Here the duplicated genes shared between the *Vac* chr2p and chrZ are likely products of whole genome duplication dated to the vertebrate ancestor, commonly referred to as ohnologs. Indeed, among all duplicated gene pairs on the chr2p and chrZ, 67% of them have been previously annotated as ohnologs (Singh et al. 2015).

By contrast, the non-repetitive homologous sequences between chr2 and chrW are only found at the 2q-end, and are concentrated at the last 20Mb region (**Figure 3c**). The RTE-BovB elements but not other TEs are also significantly (*P* < 0.05, Wilcoxon test) enriched within this 20Mb region. These results suggest that there could be one large segmental duplication (the last 20Mb region) mediated by and inserted with the RTE-BovB elements, followed by other individual duplications elsewhere in the 2q-end. We found 274 genes of chr2 with one or more duplicated copies on the chrW, 90% of which are located within the 2q-end. If gene duplications between chr2q and chrW were mediated by the varanid burst of RTE-BovBs, we expected to find enrichment of this TE family nearby the duplicated genes. Indeed, among all subfamilies of RTE-BovB, we identified md-5_family-305 and md-5_familiy-1095 that are much more enriched nearby the paralogs between the 2q-end and chrW compared to other single copy genes as a control. Other types of TEs are enriched to a similar level nearby the paralogs and the single copy genes (**Figure 3d**). In addition, we performed Transpositions in Transpositions (TinT) analyses of nested TEs, which assumes the younger TEs inserted more likely into the older ones than the opposite direction of insertion. The result supported that these two RTE-BovB subfamilies are indeed younger than other subfamilies (**Figure 3e**). Consistent with this, copies of these two subfamilies located at the 2q-end and chrW have a significantly (*P* < 0.05, Wilcoxon test) longer length, i.e., are more intact, but a significantly (*P* < 0.05, Wilcoxon test) lower divergence level from their consensus sequences, than those elsewhere in the genome (**Figure 3f**, **Supplementary Fig. 7**). Finally, when looking into the transcription of TEs across all tissues, we found only the md-5_family-1095 of RTE-BovB have pronounced expression, indicating its activity (**Figure 3g**). All these results showed that due to the no or low recombination in chrW and 2q-end regions, certain active and evolutionarily young RTE-BovB subfamilies experienced a copy number burst and preferentially accumulated at both regions in the varanid ancestor, which probably in turn mediated segmental and gene duplications between the two regions.

### Gene traffic shaped the evolution of varanid sex chromosomes

It still remains unclear whether the varanid TE burst had mediated the genes duplicated from 2q-end to chrW or the reverse. This could respectively reflect a different evolutionary force of either female-specific selection (Moghadam et al. 2012) that recruit genes elsewhere, or rescue of degenerating genes, on the chrW (Hughes et al. 2015). To discern the two scenarios, we first identified four multicopy gene families that have members located on both 2q-end and chrW in *Vac*, and also have a homolog with well annotated function in mouse. The phylogenetic trees constructed with these *Vac* gene sequences and their homologs in other non-varanid species, which experienced a lower degree of TE propagations (**Figure 3a**), thus likely have fewer or no duplications, are expected to reveal the direction of duplications into or out of chrW. These four gene families consist of *V2R*, *RNF39*, *HLA-DPB1*, and *EEF2*, with a total of 442 gene copy numbers mainly distributed on chr2 and the Z and W sex chromosomes (**Figure 4a**). All but *EEF2* have a significantly (Chi-squared test, *P* < 0.05) higher copy number in *Vac* than in three other non-varanid lizards (*Emu*, *Ppl*, and *Tiliqua scincoides* or *Tsc*). Both *RNF39* and *HLA-DPB1* are associated with immune regulation(Liu et al. 2021; Wang et al. 2021), and *V2R* or the vomeronasal chemosensory Type 2 receptor gene family encodes the pheromone receptors for environmental perception that are associated with mate choice and predator avoidance(Silva and Antunes 2017).

**Figure 4.**
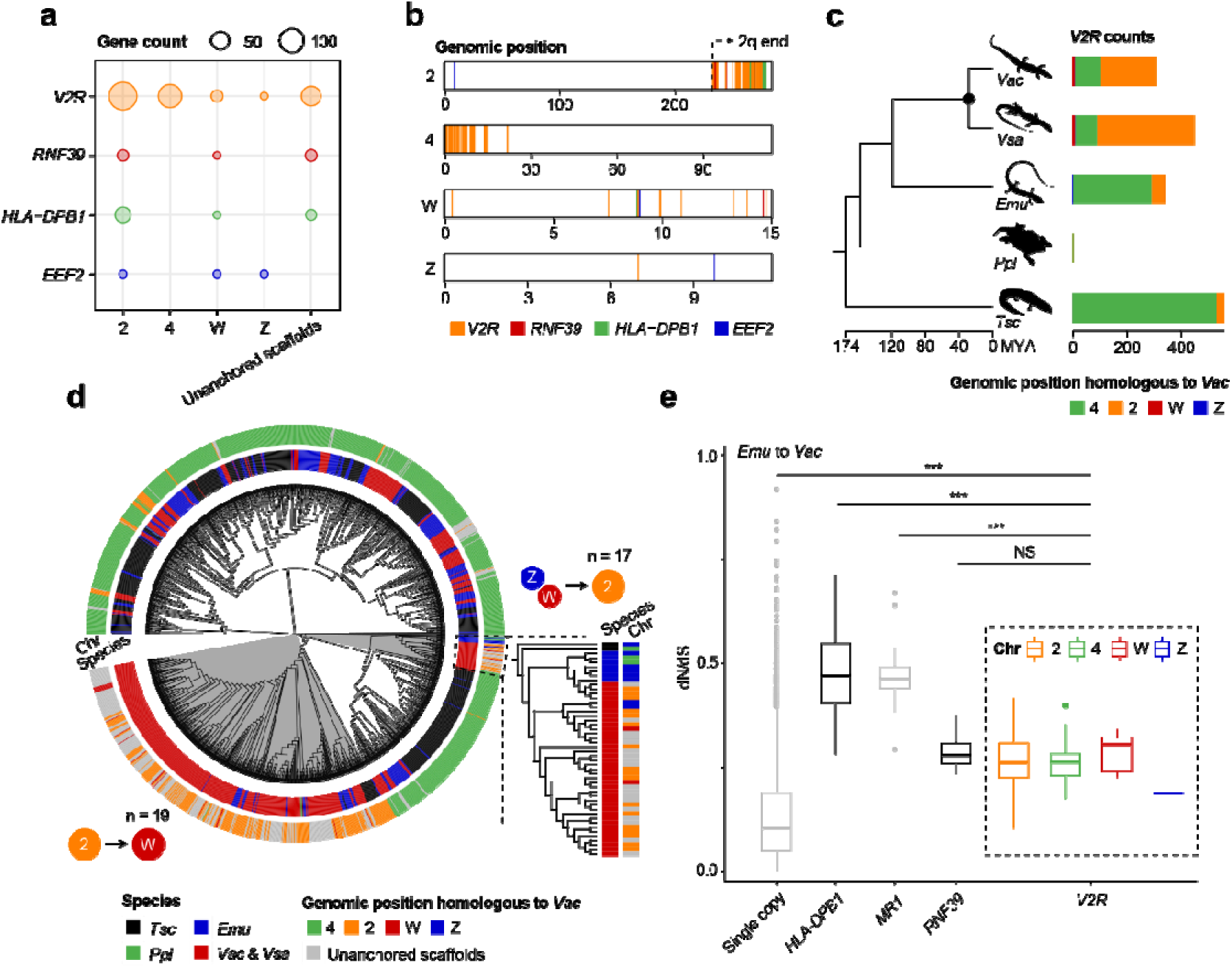
Gene traffic between the varanid chrW and chr2. **a)** Gene duplications between *Vac* Chr2 and sex chromosomes mainly belong to four gene families: *V2R*, *RNF39*, *HLA-DPB1* and *EEF2*, with the bubble size scaled to the copy number on each chromosome. **b)** Barcodes show the genomic positions of *V2R*, *RNF39*, *HLA-DPB1* and *EEF2* family members in the *Vac* genome, which are mainly concentrated at the chromosome ends of chr2 and chr4. **c)** Barplot depicted the copy numbers of *V2R* genes in several lizard species, including *Vac (V. acanthurus), Vsa (V. salvator), Emu (E. multicarinata), Ppl (P. platyrhinos), and Tsc (Tiliqua scincoides*), with different color showing the copies from different chromosomes. **d)** A maximum-likelihood tree constructed using the protein sequences of *V2Rs* from *Vac*, *Vsa*, *Emu*, *Ppl* and *Tsc* (different colors represent different species in the inner ring), with the main branches shown by different background colors as grey or white. W-linked copies (red color in the outter ring) are mainly clustered with chr2 copies (orange color). **e)** Boxplots show the pairwise dN/dS values between 1:1 orthologs in *Emu* and *Vac*, which are divided into single copy genes, the top three multicopy (copy number > 3) gene families with most abundant copy numbers throughout the genome, including *HLA-DPB1*, *MR1* and *RNF39* as a control, and *V2Rs* from different chromosomes.

*V2R*, *RNF39*, *HLA-DPB1* are mainly concentrated on the 2q-end as large tandem gene clusters, but are randomly distributed along the chrW and chrZ, with *V2R* having additional large numbers of copies on one end of chr4 (**Figure 4b**). Among the three families, the *V2R* family has probably experienced the most pronounced expansion of copy numbers in the ancestor of varanids. We annotated a total of 311 *V2R* copies throughout the genome of *Vac*, and both *Vac* and *Vsa* have about 4-fold and more than 10-fold more gene copies on the 2q-end than the outgroup species *Emu*, and even larger (> 10-fold) than in *Ppl* and *Tsc* (**Figure 4c**). While the *V2Rs* on chr4 have much less copies in the varanids than in the other lizards. Our gene trees support a different evolutionary history of *V2Rs* on different chromosomes: the *V2Rs* of chr4 form separate lineages from those of chr2 and chrW that are clustered with each other. Two lineages are of particular interests that can inform the direction of duplications. One lineage includes only the homologous *V2Rs* on the chrZ of *Emu* (whose chrW sequence is not available), those on the homologous autosome of *Tsc*, both of which are then clustered with copies from chrZ, chrW and chr2 of *Vac*. This led to an estimate of 17 *V2R* copies located on the 2q-end of *Vac* as duplication products from chrZ or chrW. While another lineage clusters the *V2Rs* of both chrW and chr2 of varanids with the chr2 copies of outgroup species, suggesting the reverse direction of duplications of 19 W-linked copies from chr2 in the *Vac* and *Vsc* (**Figure 4d**). We thus found duplications of both directions, i.e., evidence of gene traffic between the sex chromosomes and chr2 from *V2Rs*. For *HLA-DPB1* and *RNF39*, gene duplications seem to be exclusively from the 2q-end to chrW, generating only one gene copy onto the chrW. And *EEF2* probably do not have interchromosomal duplications, according to their gene trees with other lizards (**Supplementary Fig. 8**).

In particular, there are 11 *V2R* copies on the chrW, and they comprise the largest W-linked multigene family in *Vac* that is also female specific (**Supplementary Fig. 9**). As expected, 4 or 36% of these *V2Rs* have a disrupted ORF, due to either premature stop codon or frameshift mutations. This percentage is much higher than that of *V2Rs* on the autosomes, suggesting some of these W-linked copies are degenerating (**Supplementary Fig. 10**). Indeed, the evolution rates (dN/dS ratios) of *V2R* copies without an intact ORF are significantly (P < 0.05, Wilcoxon test) faster than those with an intact ORF, on both chrW and autosomes (**Supplementary Fig. 11**). Interestingly, all of the *Vsa* W-linked *V2Rs* (10 copies) have an intact ORF (**Supplementary Fig. 10**). In addition, when we compare the evolutionary rates (pairwise dN/dS ratios calculated with homologs of *Emu*) of *V2Rs* to other multicopy gene families whose copy numbers are similarly abundant in the *Vac* genome, all the *V2R* copies, including those on the chrW, show a significantly lower evolutionary rate than other multicopy gene families (**Figure 4e**), indicating some selective constraints. All these results suggest that despite being under the degenerating genetic environment of chrW, some V2R copies nevertheless have retained functions after transposition and propagation.

## Discussion

Y or W chromosome regions with complete suppression of recombination are expected to accumulate massive transposable elements and suffer long-term functional deterioration and loss of gene functions(Charlesworth and Charlesworth 2000). Therefore, chrY/W is expected to be an unfavored destination for gene transposition. In fact, independent Y-to-autosome transpositions have been reported in mammals, probably as an escape from the degenerating chrY to rescue the gene functions(Hughes et al. 2015). Contrary to this assumption, individual cases of gene acquisition on the chrY/W have been reported in *Drosophila* (Koerich et al. 2008; Carvalho et al. 2015; Tobler et al. 2017; Ricchio et al. 2021), in mammals (Page et al. 1984; Murphy et al. 2006; Li et al. 2013; Janečka et al. 2018) and in birds (Xu et al. 2020), suggesting the evolutionary course of these sex specific chromosomes can be more complex than the canonical trajectory of degeneration. Our work here indicates one of such scenarios that is mediated by a recent and local burst of TEs on an autosome, followed by directional transpositions of a large number of genes onto the chrW.

Apart from the telomeric/centromeric or nucleolar organizer regions, previous works have reported many cases of regional burst of certain TEs on one pair of autosomes (Oliveira and Wright 1998), or one chromosome of an autosome pair (Bezerra et al. 2012; Moreira et al. 2013), with the latter case often confounding the identification of heteromorphic sex chromosomes. For example, the long arm of one chr6 of platypus is enriched for satellite repeats and ribosomal genes (Rens et al. 2004). Our recent work identified some isolated populations of *Vac* and *V. citrinus* have an autosome showing cytogenetic homology to the entire chrW (Dobry et al. 2025). While over one third of the entire chr3 of several tilapia species consist of heterochromatin, which could predispose the region for becoming a sex determining region (Tao et al. 2021). Other regional repeat expansion identified in humans frequently underlie many diseases, including amyotrophic lateral sclerosis (ALS) and the fragile X syndrome (Malik et al. 2021). Although the molecular and evolutionary causes of such local TE bursts at autosomal regions are unclear, the sex specific chromosomes or other genomic regions of low recombination, like the 2q-end identified in this work, due to their inefficiency of resisting the TE transpositions, are more likely to be influenced by the burst than the rest recombining regions. When the microchromosomes or other gene-rich regions undergo such TE bursts, a byproduct of TE transposition onto the chrY/W is acquisition of a large number of genes from the autosomes like what we observed in *Vac* (**Figure 5**).

**Figure 5.**
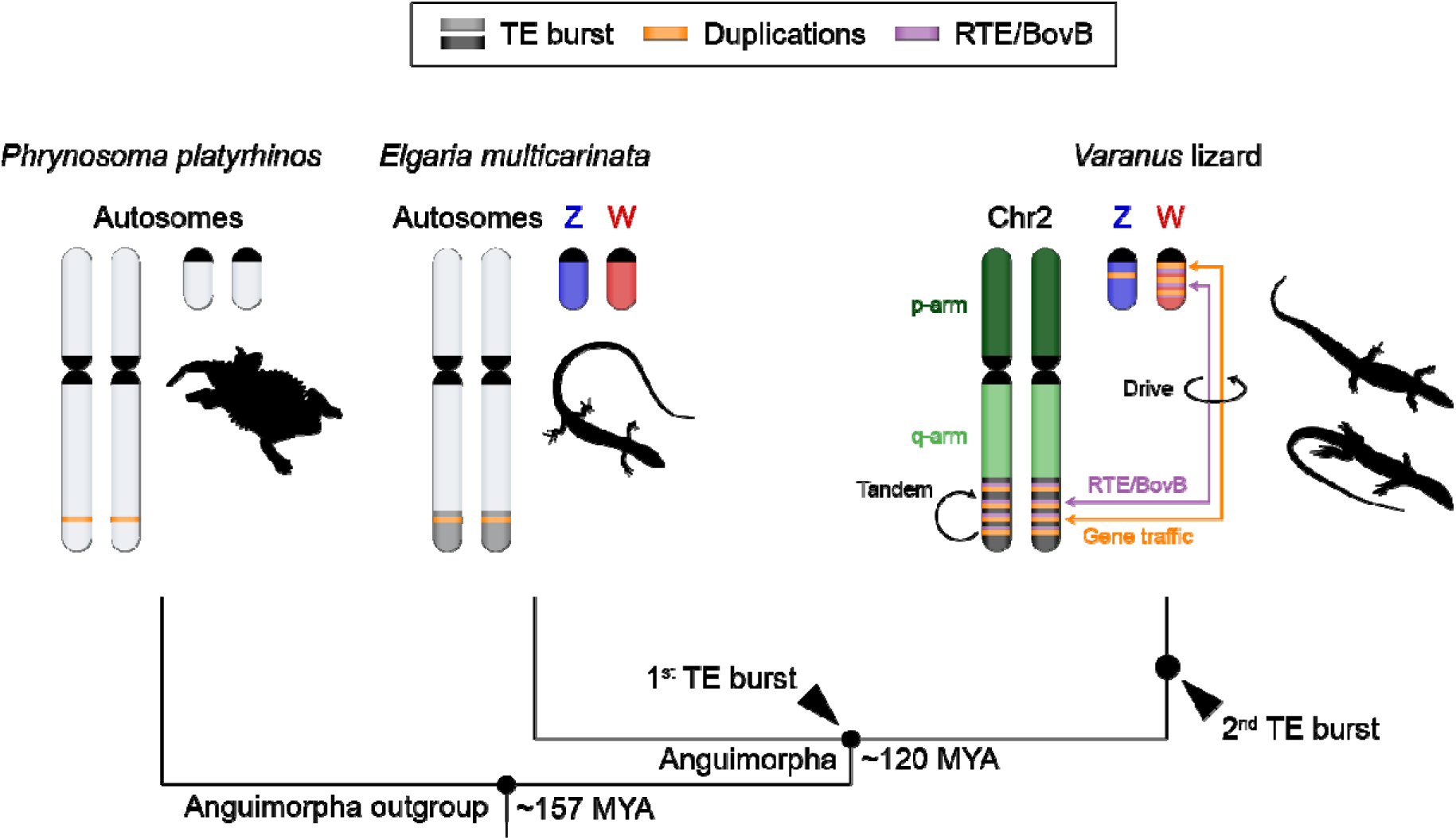
Model of gene expansion mediated by transposable elements (TEs) on sex chromosomes of *Varanus* lizards. This figure shows how consecutive expansions of TEs on autosomes during speciation of anguimorph lizards had mediated expansion of genes on their shared chrW. We inferred the first burst of TEs on the end of chr2 occurred in the Anguimorpha ancestor, and the second in the *Varanus* ancestor. The second burst has likely caused transpositions of TEs and their nearby genes between the chrW and chr2, which results in the cytogenetic homology shared only between chrW and chr2 observed in Figure 1.

The genes transposed onto the chrY/W, are then expected to undergo functional degeneration in the long-term, but at the same time, are subjected to immediate sex-specific selection. For example, in both *Drosophila* and many mammals including human, cat and cattle, some autosome-derived Y-linked single-copy gene (Carvalho et al. 2015) or multigene families (Murphy et al. 2006; Yang et al. 2011; Chang et al. 2013) have acquired testis-specific expression, in contrast to their female-biased or unbiased autosomal progenitors. Some autosome derived gene fragments were also found to be amplified on the W chromosome of a lizard *Eremias velox* (Lisachov et al. 2021) and Australian dragon lizard *Pogona vitticeps* (Ezaz et al. 2013; Matsubara et al. 2019; Alam et al. 2020). An intensively characterized example is the Y-linked *DAZ* gene family associated with human male fertility, which originated from the single copy autosomal *DAZL* gene and expanded the copy number on the Y chromosome during the primate evolution (Saxena et al. 1996; Yu et al. 2008). The example of *V2R* that we reported in this work (**Figure 5**) represents the largest autosome-to-sex chromosome transposition known to date. A previous study of the draft genome of another varanid lizard Komodo dragon (*Varanus komodoensis*) reported expansion of *V2R* (Lind et al. 2019) and here we concluded they are derived from duplications between chr2 and chrW, mediated by the TE burst. The shared large number of intact W-linked *V2Rs* between varanids during the 28.2 million years of evolution, and the pattern of evolutionary rates suggest many of them have retained functions (**Figure 4**), and possibly contribute to the sexually dimorphic perception of environmental cues in *Vac*. However, further investigation is required to demonstrate their detailed functionality, particularly tissue specific expression profile in those organs associated with expression of *V2Rs*, including the vomeronasal organ.

## Methods

### Sample collection

Individuals used in the study were collected under authorization of the University of Canberra Animal Ethics committee (Project ID: 20180306), and collection permits were obtained from the Northern Territory (Permit number 63414) and Queensland (Permit number WA0010049) governments for scientific collection. They were imported into the ACT under an import license (LT201829). All methods reported in this study were conducted in accordance with the relevant guidelines and regulations. The study is reported in accordance with ARRIVE guidelines. Collection details have been described by Dobry et al, in previous studies (Dobry et al. 2023a; Dobry et al. 2023b). Individuals were air freighted to the Canberra Airport, transported to the University of Canberra, and housed in terrariums as described by Retes and Bennett (Daniel and Frank 2001). Live tail tissue was used to culture cells for cytogenetic analysis as described previously (Dobry et al. 2023a; Dobry et al. 2023b). Euthanasia was carried out using intraperitoneal injection of sodium pentobarbitone at 60mg/kg sodium pentobarbital. Tissues were immediately collected and snap frozen with liquid N_2_ and then processed for transcriptome and DNA extraction.

### Cytogenetic works

Animal collection, microdissection, preparation of sex chromosome specific probes and validation of probes were described in our previous studies (Dobry et al. 2023b; Dobry et al. 2025). FISH mapping of sex chromosome probes are described in (Dobry et al. 2025). Briefly, sex chromosome probe sequences composed of short, labelled polynucleotides ranging from 45-47 nucleotides in length were custom designed and manufactured by Arbor Biosciences (myTags, Arbor Biosciences, Ann Arbor, MI, USA). These probes were designed from single copy regions of the sex specific chromosome scaffolds identified in a previous study (Zhu et al. 2022) and validated in (Dobry et al. 2025). Probes were used at a concentration of 100 ng/μL, with a total of 200 ng per slide with 38 μL hybridization buffer (BioCare Medical, Pacheco, CA). Coverslips were placed on each slide and sealed with rubber cement to avoid dehydration during denaturing at 68 °C for 5 min and then incubated at 37 °C for 48 h. The coverslips were then removed and the slides were washed with 0.4x SSC (3 M NaCl, 0.3 M sodium citrate, pH 7, and 0.3% (v/v) IGEPAL (Sigma-Aldrich)) at 60 °C for 2 min. A second wash was then performed with 2x SSC for 1 min. The slides were then dehydrated with a series 1 min ethanol washes consisting of 70%, 90%, and 100% (v/v) respectively and allowed to completely dry before staining with Vectashield antifade mounting medium with DAPI (Vector Laboratories). Slides were viewed and photographed with a Leica Microsystems Thunder Imaging system, and then karyotype images were constructed using Adobe Photoshop 2023.

### Genome assembly and annotation

We sequenced and assembled the reference genome of a female *V. acanthurus*. Liver tissues were collected for DNA extraction and library preparation for Illumina, PacBio SMRT, and Hi-C sequencing (See **Supplementary Table S1**). Original assembly was performed with Flye (v2.7-b1585) (Kolmogorov et al. 2019) using PacBio reads and polished with Illumina DNA reads using Pilon (v1.22) (Walker et al. 2014). The assembly was subsequently scaffolded with Ragtag (v2.1.0) (Alonge et al. 2022) using BGI stLFR reads, and completed with the 3D-DNA pipeline (v180922) (Dudchenko et al. 2017) based on Hi-C data, resulting in an assembly with over 97% of sequences assigned to chromosomes, and BlobToolKit (v4.3.10) (Challis et al. 2020) were later used for genome quality control. For genome annotation, a de novo repeat sequence database was first constructed using RepeatModeler (v2.0.3) (Flynn et al. 2020), and combined with the squamate repeat consensus library from Repbase (v20181026) (Bao et al. 2015) to create a comprehensive repeat library. Repeat sequences in the genome were then identified and classified with RepeatMasker(Smit et al.) (v4.07). Later we used the Funannotate pipeline (v1.8.5) (Palmer and Stajich) for the gene annotation, obtaining transcriptome data from brain and gonad tissues, each with two replicates per sex for comprehensive transcript assembly. To construct a non-redundant reference protein library, we included proteins from Komodo dragon (*Varanus komodoensis,* GCF_004798865.1, (Lind et al. 2019)), green anole (*Anolis carolinensis,* GCF_000090745.2, (Alföldi et al. 2011)), Indian cobra (*Naja naja,* GCA_009733165.1, (Suryamohan et al. 2020)), chicken (*Gallus gallus,* GCF_016699485.2, (Rhie et al. 2021)), mouse (*Mus musculus,* GCF_000001635.27, (Church et al. 2011)), and human (*Homo sapiens,* GCF_000001405.40, (Schneider et al. 2017)) using cd-hit(v4.7) (Fu et al. 2012). Among all annotated genes, those with transcripts containing more than two premature stop codons or frameshift mutations were classified as having disrupted open reading frames (ORFs), and genes with more than two copies were classified as multicopy genes.

### Sex chromosome identification

To identify the sex chromosome sequences of *V. acanthurus*, Illumina DNA reads from both sexes were aligned to the masked genome using Bowtie2 (v2.2.9) (Langmead et al. 2019). Read coverage for each sex was calculated in 100-kb non-overlapping windows using SAMtools (v1.6) (Danecek et al. 2021) and normalized to the median depth per base pair across the entire genome. This normalization facilitated direct comparison of coverage between sexes. The "covered site" was defined as a base pair with a read coverage of at least 3. The Z chromosome was expected to display an autosome-like male-to-female (M/F) coverage ratio but with half the overall read coverage due to hemizygosity in females. Conversely, the highly differentiated W chromosome was identified by scaffolds exhibiting both an M/F coverage ratio and a covered site ratio below 0.2, alongside a scaffold size exceeding 5 kb. These W-linked scaffolds were subsequently assembled into a contiguous W chromosome using Ragtag (v2.1.0), with the Z chromosome serving as a reference (**Supplementary Table S4**). For *V. salvator*, where Illumina DNA reads (SRR16080542) were available only from female tissues, autosomes and sex-linked sequences were distinguished based on sequencing depth. Autosomal sequences displayed approximately two-fold the depth of sex-linked sequences. Homology-based comparisons with the Z and W chromosomes of *V. acanthurus* were then employed to classify W-linked and Z-linked sequences in *V. salvator*.

### Evolutionary strata

To identify the evolutionary strata of the sex chromosomes in *Varanus acanthurus*, we first identified regions with similar sequencing depth between sexes, as the pseudoautosomal region (PAR). The sequence similarity between the Z and W chromosomes was then assessed by aligning the masked W chromosome sequence to the Z chromosome using Lastz (v1.02.00) (Harris 2008) with parameters “--hspthresh=2200 --inner=2000 --ydrop=3400 -- gappedthresh=10000”. Syntenic fragments were merged into longer blocks through the alignment chains and nets by the UCSC pipeline. Sequence similarity between the sex chromosomes was measured in 100-kb sliding windows along the Z chromosome. To evaluate male heterozygosity along the Z chromosome, SNP calling was performed using the HaplotypeCaller module from the Genome Analysis Toolkit (GATK, v3.8) (Van der Auwera and O’Connor) with male Illumina DNA reads. SNPs were filtered under the following criteria: "QD < 2.0, MQ < 40.0, FS > 60.0, SOR > 3.0, MQRankSum < −12.5, and ReadPosRankSum < −8.0." SNP counts for each chromosome were calculated in 100-kb windows. Both sequence similarity and male heterozygosity values were used to change-point analysis using the R package cpm (v2.3) (Ross 2015). The resulting strata were statistically evaluated using the Wilcoxon-test to confirm significant differences between strata.

### Gene expressions

To quantify the gene expressions, RNA-seq reads from brain tissues of *Varanus acanthurus* of both sexes, as well as testis and ovary, were mapped to the genome using Hisat2 (v2.1.0) (Kim et al. 2019), retaining only uniquely mapped reads (flagged as NH:i:1 and without the ZS:i flag). FeatureCounts (v1.6.2) (Liao et al. 2014) were used to count these reads, and gene expression levels were normalized and quantified as TPM (Transcripts Per Million). A similar pipeline was employed to quantify gene expression levels in chicken, utilizing RNA-seq reads from liver and muscle tissues of both sexes, as well as testis and ovary (see **Supplementary Table S2**). For all figures, the expression levels of each gene in each tissue were represented by the median values across replicates. Genes with TPM < 1 across all tissues were classified as silenced.

### Inferring synteny between genomes

To identify synteny blocks between species in this study, reciprocal best-hit (RBH) blast was performed using BLASTp (v2.6.0) to detect one-to-one orthologs based on the protein sequences of all species. The search employed a stringent e-value threshold of 1e-10, and results were filtered to retain alignments covering more than 50% of the reference protein and exhibiting at least 50% sequence identity. In addition to *Varanus acanthurus*, the species included in this comparative study were another monitor lizard (*Varanus salvator*, NCBI accession JAIXND000000000.1, (Chetruengchai et al. 2022)), a lizard from Anguimorpha (*Elgaria multicarinata*, GCF_023053635.1), a lizard from Iguania (*Phrynosoma platyrhinos*, GCA_020142125.1, (Koochekian et al. 2022)), a skink (*Tiliqua scincoides*, GCF_035046505.1, (Rhie et al. 2021)), and chicken (*Gallus gallus*, GCF_016699485.2, (Rhie et al. 2021)).

### dN/dS calculation

Pairwise dN/dS values between orthologs were calculated as follows: one-to-one orthologs were first identified using the RBH pipeline described earlier. Coding sequences for each ortholog pair were aligned using Prank (v150803) (Löytynoja 2014) with default parameters. dN/dS values were then calculated for each pairwise alignment using the *codeml* module of PAML (v4.8) (Yang 2007)under the free-ratio model. The same pipeline, with identical parameters and modes, was applied to analyze orthologs between *Varanus acanthurus* and *Elgaria multicarinata*, as well as *Varanus acanthurus* and *Phrynosoma platyrhinos*.

### Repeat analysis

To assess the relative abundance of retroposons at the distal ends of chromosomes 2 and the whole W in varanids compared to homologous genomic regions in the other species, we calculated the length of each repeat family in 100kb windows across these chromosomes and normalized by the chromosome-wide mean. To investigate the specific repeat subfamilies potentially driving gene duplication events, repeats located within ±10kb of a gene were categorized as nearby repeats. To quantify enrichment, we calculated the proportion of genes with nearby repeats from each subfamily, along with the proportion of copy number of these nearby repeats relative to total loci on the chromosomes. To assess the relative activity and age of retroposons, we first applied TinT (Transpositions in Transpositions) algorism to examine the frequencies of nested transpositions. We also used RepeatMasker outputs to calculate Kimura distances for each retroposon subfamily to measure the divergence level of each element. For further phylogenetic inference, consensus sequences of the RTE/BovB subfamilies from two varanids and the desert horned lizard (*Ppl*) were analyzed using IQ-TREE.

RNA-seq data from brain tissues of both sexes, male testis, and female ovary were mapped to the genome using the same pipeline as employed for gene expression analysis. The resulting BAM files were subsequently processed with TEcount (v1.0.1) (Marasca et al. 2022) for quantification of counts by class, family and subfamily levels. Repeat expression levels were later normalized as CPM (Counts Per Million). For all figures, the expression levels in each tissue were represented by the median values across replicates.

### Paralog analysis

To first identify paralogs in the genome of each species, the longest transcript of each gene in *Vac*, *Vsa*, *Emu*, *Ppl* and *Tsc* were fed to OrthoFinder (v2.5.4) (Emms and Kelly 2019) with default parameters, to get orthogroups that pinpoint inter- and intra-species duplications. For the identification of ohnologs, the defined duplications were queried against the database at http://ohnologs.curie.fr/, using the chicken data as a reference. Later, protein sequences of all paralogs within each orthogroup were first aligned using MAFFT (v7.505) (Katoh and Standley 2013) with default parameters, and poorly aligned regions were trimmed with TrimAl (v1.5.rev0) (Capella-Gutiérrez et al. 2009) with -gt 0.1 option. Maximum likelihood trees were then inferred using IQ-TREE (v1.6.11) (Minh et al. 2020). The direction of the duplication events in varanids were inferred based on both the genomic positions of their homologs in the outgroup species *Emu*, *Ppl* and *Tsc,* and the structure of phylogeny tree from each gene family.

### Data Access

The genome assembly is available on NCBI under accession numberPRJNA855548. The annotation file is available on Github: https://github.com/zjuzexian/Varanus-sex-chromosomes/. Sequenced reads generated during this study are available on NCBI PRJNA1201623. The stLFR reads, produced in our previous study, can be accessed under accession ID PRJNA737594. *Varanus acanthurus* draft genome assembly that are used to develop sex chromosome specific probes are available via genome warehouse PRJCA005583 and sex chromosome probe sequences are available in Dobry et al (Dobry et al. 2025). Note the codes used to generate the figures can be found on GitHub: https://github.com/zjuzexian/Varanus-sex-chromosomes/

## Acknowledgement

Qi Zhou is supported by the National Key Research and Development Program of China (2023YFA1800500), National Natural Science Foundation of China (32170415). Jason Dobry is partially supported by the Australian Government Research Training Program (RTP) stipend scholarship. Tariq Ezaz and Erik Wapstra were partially supported by the Australian Research Counil Discovery Project grant (ARC DP200101406).

